# Infra-slow modulation of fast beta/gamma oscillations in the mouse visual system

**DOI:** 10.1101/2020.02.24.963124

**Authors:** Patrycja Orlowska-Feuer, Annette Elisabeth Allen, Timothy Matthew Brown, Hanna Jowita Szkudlarek, Robert James Lucas, Riccardo Storchi

## Abstract

Infra-slow (<0.02 Hz) and fast beta/gamma (20 – 100 Hz) oscillations in neurophysiological activity have been widely found in the subcortical visual system. While it is well established that fast beta/gamma oscillations are involved in visual processing, the role (if any) of infra-slow oscillations is currently unknown. One possibility is that infra-slow oscillations exert influence by modulating the amplitude of fast oscillations, yet the extent to which these different oscillations arise independently and interact remains unknown. We addressed these questions by recording *in vivo* spontaneous activity from subcortical visual system of visually intact mice, and animals whose retinal network was disrupted by advanced rod/cone degeneration (*rd/rd cl*) or melanopsin loss *(Opn4^-/-^* We found many neurons expressing only one type of oscillation, and indeed fast oscillations were absent in *rd/rd cl.* Conversely, neurons co-expressing the two oscillations were also common, and were encountered more often than expected by chance in visually intact but not *Opn4^-/-^* mice. Finally, where they co-occurred we found that beta/gamma amplitude was modulated by the infra-slow rhythm. Our data thus reveal that: 1.) infra-slow and beta-gamma oscillations are separable phenomena; and 2.) that they actively co-occur in a subset of neurones in which the phase of infra-slow oscillations define beta-gamma oscillation amplitude. These findings suggest that infra-slow oscillations could influence vision by modulating beta-gamma oscillations, and raise the possibility that disruptions in these oscillatory behaviours contribute to vision dysfunction in retinal dystrophy.

**KEY POINTS SUMMARY:**

- Neurophysiological activity in the subcortical visual system fluctuates in both infra-slow and fast oscillatory ranges, however the level of co-occurrence and potential functional interaction of these rhythms is unknown.
- Analyzing dark-adapted spontaneous activity in the mouse subcortical visual system, we find that these two types of oscillation interact uniquely through a population of neurons expressing both rhythms.
- Genetic ablation of rod/cone signaling potentiates infra-slow and abolishes fast beta/gamma oscillations while genetic ablation of melanopsin substantially diminishes the interaction between these two rhythms.
- Our results indicate that in an intact visual system the phase of infra-slow modulates fast beta/gamma oscillations.
- Thus one possible impact of infra-slow oscillations in vision is to guide visual processing by interacting with fast narrowband oscillations.

## INTRODUCTION

Regular infra-slow (<0.02 Hz) oscillations in neurophysiological activity have been found throughout the mammalian visual system, including the retina (Rodieck & Smith, 1966), optic chiasm (Cavaggioni, 1968), the dorsal lateral geniculate nucleus (dLGN) (Albrecht & Gabriel, 1994; Albrecht *et al*., 1998), ventral lateral geniculate nucleus (vLGN) (Chrobok *et al*., 2018), olivary pretectal nucleus (OPN) (Szkudlarek *et al*., 2008, 2012), suprachiasmatic nucleus (SCN) (Aggelopoulos & Meissl, 2000), intergeniculate leaflet (IGL) (Lewandowski *et al.*, 2000, 2002) and primary visual cortex (Filippov & Frolov, 2005). Infra-slow oscillatory rhythm features such as frequency and amplitude are modulated by light and depend on retinal activity (Miller & Fuller, 1992; Albrecht *et al*., 1998; Lewandowski *et al*., 2000; Szkudlarek *et al*., 2012; Orlowska-Feuer et al., 2016a). Nevertheless, despite its ubiquity, the contribution of such rhythmicity, if any, to visual processing is still a matter of debate.

Oscillations at a quite different frequency range (20 – 100 Hz, termed beta/gamma) are also a feature of the early visual system. Such oscillations have been implicated in different aspects of visual processing ranging from inter-area communication to feature binding (Gray *et al.*, 1989; Fries, 2005). Recently, we and others also reported that the power of fast beta/gamma oscillations encodes for ambient light intensity and spatially structured changes in luminance (Saleem et al., 2017; Storchi et al., 2017).

Decades of research have highlighted a general organizing principle by which the brain orchestrates oscillations expressed at different timescales. Namely, faster oscillations are “nested” within slower oscillations so that the power of faster oscillations is controlled by the phase of the slower ones. Remarkable example of this phenomenon is the theta-gamma coupling in the hippocampus that has been shown to underlie spatial navigation and object memory recollection (Lisman & Jensen, 2013; Gupta et al., 2016). Electroencephalographic recordings in humans indicates that also infra-slow oscillations can modulate faster rhythms (Vanhatalo *et al*., 2004; Monto *et al*., 2008). Thus one possible impact of infra-slow oscillations in vision is to guide visual processing by interacting with fast narrowband oscillations.

To test this possibility we recorded from mouse subcortical visual system (dLGN, vLGN, and OPN). We found that infra-slow and fast beta/gamma oscillations interact through a population of neurons that co-express both rhythms. Namely in these neurons the phase of infra-slow modulates the amplitude of gamma oscillations. We then asked whether this interaction required intact vision by repeating the same experiments in animals with genetic ablation of rods+cones or melanopsin photoreception. We found that rod+cone loss abolished fast oscillations, but left infra-slow rhythm intact, whereas both oscillations were retained in melanopsin knockouts but their co-expression was substantially reduced. Together these results confirm that infra-slow and fast oscillations are separable phenomena reflecting different network states, and also that in physiological conditions their co-expression in the same neurons is an active consequence of a particular network state that is disrupted by melanopsin loss.

## MATERIALS AND METHODS

### Ethical approval

All procedures involving animals were carried out in accordance with regulations and standards of the European Community Council Directive of 24 November 1986 (86/609/EEC), approvals of the institutional ethics committee and the UK Home Office and standard and regulations described by (Grundy, 2015).

The following report is in part a re-analysis of previously collected and partially published elsewhere data (Brown *et al*., 2010; Allen *et al*., 2011).

### Animals

Experiments were performed on genetically modified mice which were bred in the animal facility of the Faculty of Life Sciences at Manchester University, UK. Animals were housed under standard conditions (60% of humidity and food and water *at libitum)* with 12:12 h light/dark cycle. Animals were 3 – 5 months old males with three different genetic backgrounds: *Opn1mw^R^* mice (C57BL/6:129sv strain) had all types of photoreceptors functional; rodless and coneless *(rd/rd cl)* mice (C3H strain) lacked rods and cones and *Opn4^-/-^* mice (C57BL/6:129sv strain) lacked melanopsin. *Opn1mw^R^* mice had the medium wavelength-sensitivity (MWS) opsin replaced by human red opsin. This manipulation shifts cone spectral sensitivity but has no other known effects on retinal functions and connectivity (Smallwood *et al*., 2003; Jacobs & Williams, 2007). Therefore, since the manipulation is inconsequential for this study, *Opn1mw^R^* mice were used as ‘wild type’ animals. All animals were removed from their home cages at the beginning of the projected day and the electrophysiological recordings spanned the mid portion of the day. In total 61 animals were used: 22 of *Opn1mw^R^*, 22 of *Opn4^-/-^* and 17 of *rd/rd cl* mice.

### Surgery

All surgical procedures were conducted under deep urethane anesthesia (1.55 g / kg, 30% w / v; Sigma-Aldrich, Munich, Germany). Animals were injected intraperitoneally and the depth of anesthesia was ascertained by lack of withdrawal and ocular reflexes. If necessary, animals were supplemented with 10% of the initial dose of urethane, however that has never been done during the recordings. Body temperature was automatically maintained at 37 ± 0.5°C by the thermistor with a feedback-controlled heating pad. Anesthetized mice were carefully placed in a stereotaxic frame via ear bars (SR-15M, Narishige International Limited, London, UK). Skin and all soft tissues covering the skull were removed exposing coronal, sagittal and lambdoid sutures. Next, bregma point was determined and craniotomy was performed above the OPN or LGN. The coordinates were assessed based on a stereotaxic brain atlas for mice (Paxinos & Franklin, 2001) and were AP, −2.8; LM, 0.9; DV, −2 to −2.8 mm and AP, −2.5; LM, 2.2; DV, 2.5 to 3.5 mm for the OPN and LGN, respectively.

### Electrophysiology

A 32-channel multiunit electrode array (A4X8-5 mm-50-200-413; NeuroNexus Technologies, Dallas, TX, USA) was used to record neuronal activities from the pretectum and LGN area. It consisted of 4 silicon shanks (spaced 200 μm), each with 8 recordings sites spaced vertically at 50 μm. Neuronal signals were acquired using Recorder64 system (Plexon, Dallas, TX, USA). Signals were amplified (3000×), filtered (300 Hz) and digitized at 40 kHz. All data were saved on a computer hard disc for further analysis.

A 4 × 8 electrode array was introduced to a 350 × 600 μm area of the pretectum or LGN and recordings were taken regardless of types of recorded activities (either oscillatory or non-oscillatory). All experiments were conducted in dark adapted conditions provided by coverage of the Faraday cage with light-impermeable material (effective photons flux ~ 9 log photons cm^-2^ s^-1^). Moreover, recordings in some animals were not limited to one penetration.

### Histological verification

Prior to the first insertion, the electrode was dipped in fluorescent dye (Cell Tracker CM-Dil, Invitrogen Ltd., UK) in order to verify position of all recording places. At the end of the experiment animals were intracardially perfused with a buffered physiological saline followed by 4% paraformaldehyde in 0.1 M phosphate buffer (pH = 7.2). Brains were removed from the skull and postfixed overnight in the same fixative at 4°C. Cryoprotected brains (immersion in 30% sucrose solution for 2-3 days) were cut on a freezing microtome at a coronal plane. Sequential sections (100 μm thick) with CM-Dil depositions were mounted on the gelatin-coated glass slides. Sections were carefully inspected, photographed and ascribed to a related coronal plain of the brain according to the stereotaxic atlas of a mouse brain (Paxinos & Franklin, 2001). Expression of the parvalbumin (calcium binding protein) was used as the marker of OPN neurons (for the precise description of the method see Allen et al., 2011). The optic tract, hippocampus and third ventricle were used as landmarks to determine the LGN borders.

### Statistics

#### Spike Sorting

Multi-unit recordings were processed using Offline Sorter (version 2.8.8; Plexon) or Spike2 (version 6.08). After removal of cross-channel artifacts, each channel was analyzed separately. Single-units were detected and categorized based on spike waveform via principal component analysis (PCA) and related statistic (Fig. 2A-B). Moreover, the inter-spike interval (ISI) histograms were computed to monitor unit refractory period. Cross-correlograms for all isolated units were examined to ensure that each unit was only included once in the further analysis (probability of synchronous firing <0.01). In total we detected 2606 neurons and 2372 (those with firing rates >0.02Hz) were further analysed.

#### Detection of Infra-Slow activity

Regular oscillations in the infra-slow range has been previously shown in several publications (Miller & Fuller, 1992; Albrecht & Gabriel, 1994; Lewandowski *et al*., 2000; Szkudlarek *et al*., 2008; Orlowska-Feuer *et al*., 2016a; Tsuji *et al*., 2016; Chrobok *et al*., 2018). The frequency of those oscillations has been shown to be <0.02 Hz even when long recordings epochs (>1hour, (Orlowska-Feuer *et al*., 2016a; Chrobok *et al*., 2018)) where considered. However in such works the assessment has been largely done qualitatively based on visual inspection of firing rate time series, autocorrelations and power spectra. Here we devised a procedure to classify those units automatically. We first observed that, for units expressing the most regular infraslow oscillations, their autocorrelation could be well fit by using the following equation:

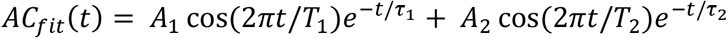

Where *e*^-*t*/*τ_j_*^ captures the reduction in autocorrelation as function of time lag *t* and cos(2*πt*/*T_i_*) captures regular oscillations (Fig. 1A-B). The dominant oscillation was defined so that for all units *A*_1_*τ*_1_ > *A*_2_*τ*_2_. The ratio *T*_1_/*T*_2_ clustered around 0.5 and 2 indicating that the second oscillation typically approximates a harmonic of the dominant oscillation (Fig. 1C). For classification we focussed on the dominant oscillation. In order to classify a unit as infra-slow we used the following criteria:

1. Consistent with previous literature the period T1 has to be within the 50s and 450s (Miller & Fuller, 1992; Lewandowski *et al*., 2000; Filippov *et al*., 2004; Szkudlarek *et al*., 2008)
2. The relation *τ*_1_/*T*_1_ between time constant and oscillation period has to be above a fixed threshold (=0.6) in order to reveal regular oscillations (see Fig. 1D-F).
3. The autocorrelation model of equation 1 has to reduce (by >22.5%) the relative fitting error over a simpler model described by *AC_fit simple_*(*t*) = *Ae*^-*t/τ*^.

**Fig. 1.**
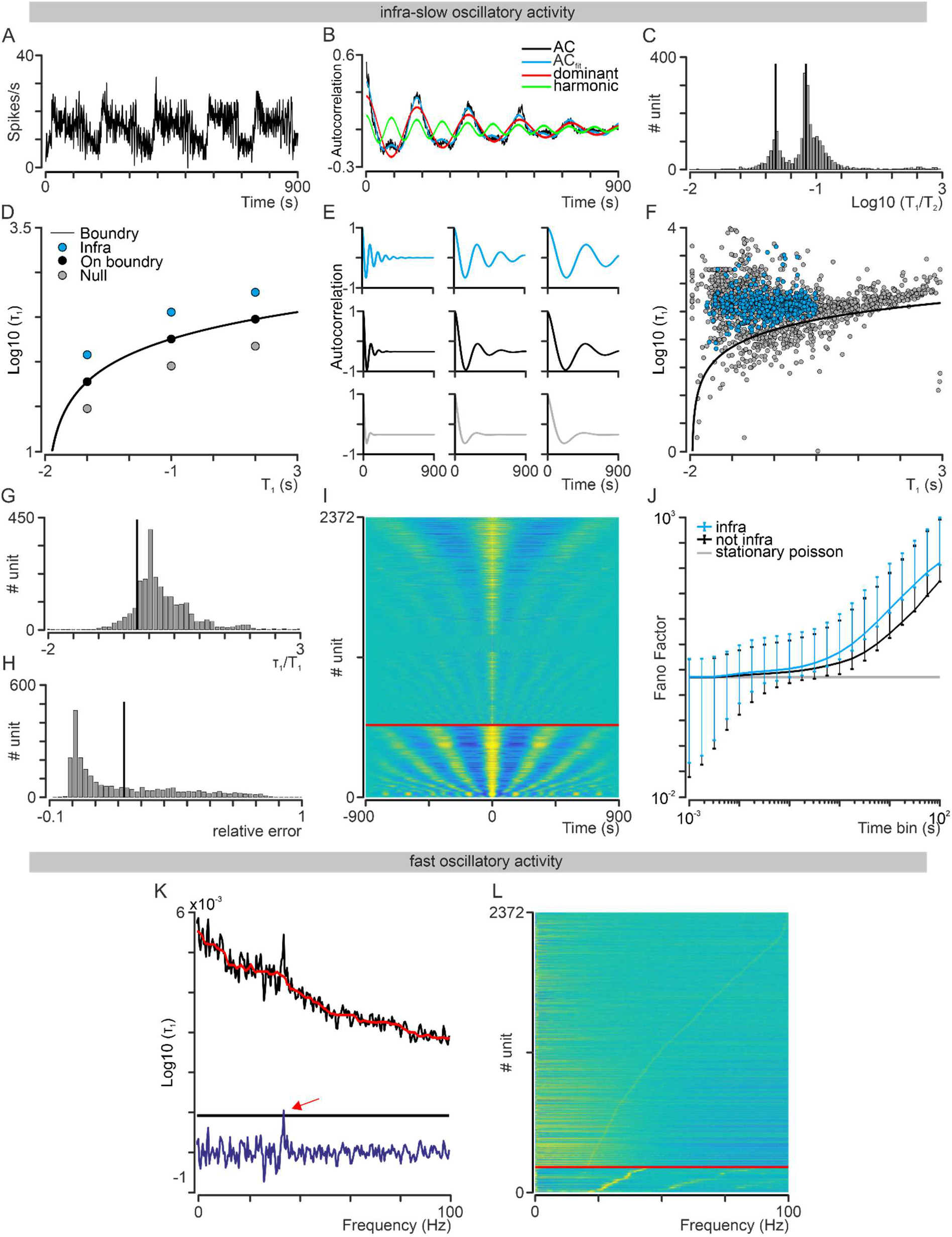
Infra-slow and fast beta/gamma criteria for classification. A) Firing pattern of a representative unit expressing regular infra-slow oscillations. B) The autocorrelogram (black) for unit in panel A can be fit with a simple model expressed in **equation 1** (blue) and composed of a dominant (red) and a sub-dominat (harmonic, green) component. C) Histogram of the ratio between period of the dominant and sub-dominant component in **equation 1** for the whole dataset (n = 2372). D) Visual representation of the second classification criterion for classifying unit as infra-slow: the ratio τ_1_/*T*_1_ between time constant and oscillation period has to be above a fixed threshold (= 0.6). Blue dots represent surrogate units meeting the criterion, grey dots represent units that don’t meet the criterion and black dots units located on the boundary. E) Autocorrelogram for the units in panel D. F) Same as panel D for all the units in the dataset (n = 2372). Blue dots indicate units that meet all the three criteria for infra-slow classification. G, H) Histograms for *τ*_1_/*T*_1_ ratio (G) and relative error (H) for all units in the dataset (n = 2372). I) Crosscorrelogram for all units in the dataset (n = 2372). Units classified as infra-slow are displayed at the bottom below the red line. All units are re-ordered according to *T*_1_ parameter. J) Fano-factor as function of time window for spike counts for infra-slow units (blue; mean & 0.01-0.99 confidence interval) and for units that do not express regular infra-slow oscillations (black; mean mean & 0.01-0.99 confidence interval). Both groups deviate from a stationary Poisson process (grey line). K) Detection of narrowband oscillations in the beta/gamma range. The PSD for a representative unit (black) is rectified by using a median filter (red). A threshold of 5SD is the applied to the rectified PSD (blue) to detect the narrowband peak. L) PSD for all units in the dataset (n = 2372). Units classified as expressing fast beta/gamma oscillations are displayed at the bottom below the red line. All units are re-ordered according to oscillation frequency (note the narrowband peaks indicated by the red arrow).

Since no clear indication of bimodality was present in the data (see Fig. 1G-H) numerical thresholds for Criteria 2 & 3 were derived empirically to match manual qualitative classification on a subset of data. However the full dataset clearly shows a neat separation between units that were classified as infra-slow and those that were not (Fig. 1I). Consistently with results obtained in cat LGN by (Teich *et al*., 1997) both units classified as infra-slow and those that were not exhibited fluctuations in the infra-slow range (Fig. 1J).

#### Detection of Fast Beta/Gamma activity

We first calculated the Power Spectral Density (PSD) for each unit (Fig. 1K) by using the Welch method (time bin = 1s across for whole 900s recording epoch). For this analysis we applied Hamming windows to 512s epochs and 50% overlap between epochs (see MATLAB function *pwelch.m).* In order to identify narrowband peaks in the beta/gamma range we de-trended the PSD by using a one dimensional median filter (Fig. 1K; filter order = 20; see MATLAB function *medfilt1.m).* The threshold for a narrowband peaks was then calculated as equalling 5 times the standard deviation of the de-trended PSD (Fig. 1K). The units classified as expressing beta/gamma oscillations showed a clear narrow peak in PSD (Fig. 1L).

#### Estimation of Cross-Correlation Index

A cross-correlation function was estimated between all pairs of units simultaneously recorded. For each pair a cross-correlation index was then obtained by calculating the median of the cross-correlation function in the interval (−10, 10) seconds. To test whether this index was significantly different from zero we applied a sign-test at the distribution of cross-correlation indexes across the selected units.

#### Estimation of coupling between infra-slow and fast beta/gamma oscillations

Spike timestamps were binned at 1 s for infra-slow and 5 ms for fast beta/gamma to obtain spike counts. Those counts were filtered by using zero-phase Kaiser filters (infra-slow: low-pass filter with cut-off at 2.5/*T*_1_ Hz and stop-band at 5/*T*_1_ Hz; fast beta/gamma: band-pass filter with cut-off at 0.8 and 1.2 times the peak frequency and stop-band at 0.6 and 1.4 times). A Hilbert transformation was then applied to the spike counts filtered in the infra-slow range and the phase of these signals was clipped in the range (-π, π). Finally the phase of infra-slow spike counts was used to divide fast beta/gamma band spike counts into the intervals ((0, π/2); (π/2, π); (0, −π/2); (−π/2, −π)). In order to test against the possibility that the amplitude of fast beta/gamma oscillations is modulated by the phase of infra-slow oscillations we compared the amplitude of these oscillations in the range (-π/2, π/2) with the amplitude observed in the complementary range by using a non-parametric sign-test.

#### Permutation Test

Permutation test was used to test the null hypothesis that occurrence of fast and infra-slow oscillations at the level of individual units were statistically independent events. First we counted the actual number of units expressing both fast and infra-slow rhythms. Then we compared those numbers with a null distribution obtained using the following simple permutation technique. First, for each recording, we generated a binary N-by-2 matrix where each row represented an individual unit, the first column reported presence (or absence) of infra-slow oscillations (presence = 1; absence = 0) and the second column presence (or absence) of fast oscillations. Then, in order to break statistical dependence between these oscillations, we performed a random permutation of one of those columns. Finally, we calculated the number of units co-expressing both rhythms (equivalent to a logical AND). Since different realization of a random permutation can produce different results we estimated the null distribution by repeating this procedure 100000 times. An example is shown in Table 1.

**Table 1:** Permutation Test. The original table contains the data where infra-slow and fast oscillatory expression was detected, the co-expression column tags the units where infra-slow and fast oscillations were co-expressed. The total number of those units is reported as TOTAL COEXP. In the permutation table a random permutation has been applied to the fast column resulting in a different number of units co-expressing both rhythms.

| Original Table |  |  |  | Permutation Table |  |  |  |
| --- | --- | --- | --- | --- | --- | --- | --- |
| #unit | Infra-slow | Fast | Co-expression | #unit | Infra-slow | Fast | Co-expression |
| #1 | 1 | 0 | 0 | #1 | 1 | 1 | 1 |
| #2 | 1 | 1 | 1 | #2 | 1 | 0 | 0 |
| #3 | 0 | 0 | 0 | #3 | 0 | 0 | 0 |
| #4 | 0 | 0 | 0 | #4 | 0 | 1 | 0 |
| #5 | 1 | 1 | 1 | #5 | 1 | 0 | 0 |
| #6 | 1 | 1 | 1 | #6 | 1 | 0 | 0 |
| #7 | 0 | 0 | 0 | #7 | 0 | 1 | 0 |
| #8 | 1 | 0 | 0 | #8 | 1 | 0 | 0 |
| #9 | 0 | 0 | 0 | #9 | 0 | 0 | 0 |
| TOTAL<br>COEXP |  |  | 3 | TOTALCOEXP |  |  | 1 |

In the real data the firing rate distribution is different between units expressing infra, fast beta/gamma or no oscillations (see Fig. 4G and Fig. 6G). Therefore, the number of units coexpressing infra and fast beta/gamma (*N_coexpress_*) expected by chance would equal:

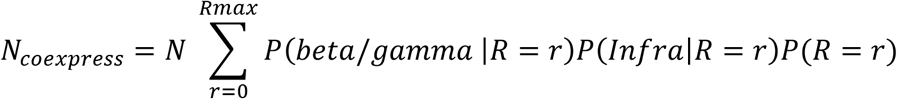

where *N* represents the total number of units and *R_max_* the maximum firing rate. Thus the random permutation procedure needs to preserve the distributions *P*(*beta/gamma\R* = *r*) and *P*(*Infra\R* = *r*). We implemented this constraint by grouping units into different classes of firing rates and permuting only across units within the same class. Moreover, in order to also exclude confounding effect of different anatomical structures, random permutations were only applied among units recorded from the same structure.

#### Statistical Comparisons

Statistical comparisons of the means ± SD were performed using non-parametrical Mann-Whitney test, Sign-test and Kruskal-Wallis test (followed by Dunn’s Multiple Comparison Test). The sampling distribution was verified with Chi-square test. A probability error of p < 0.05 was regarded as significant. All data were visualized and statistically examined using GraphPad (version 4.02; GraphPad Software), NeuroExplorer (version 5; Nex Technologies), Statistica software (StatSoft, Inc., Tulsa, Oklahoma, USA) and scripts written in MATLAB (R2019a).

#### Codes Accessibility

The original dataset in .csv format and MATLAB codes for replicating the tests are available online at https://github.com/RStorchi/InfraSlow.

## RESULTS

The main aim of this work was to determine the relationship between infra-slow and fast narrowband oscillations in the mouse visual system. We addressed these questions both in image forming and in non-image forming centres in the brain by focussing on the dLGN (image forming), vLGN and OPN (both non-image forming structures).

### Infra-slow oscillations are widespread in the mouse visual system & co-exist with fast narrowband rhythms

Infra-slow oscillations in the subcortical visual system have been observed in many species including rats, cats and monkeys but not previously in mice. Therefore, we first aimed to verify the occurrence of infra-slow oscillatory activity in the spontaneous neuronal firing in the mouse subcortical visual system. We unbiasedly recorded in the absence of visual input (in darkness) from the OPN, dLGN and vLGN in wild type mice.

Autocorrelation analysis revealed that ~30% of all recorded units expressed an infra-slow oscillatory rhythm (0.02 – 0.0022 Hz) in spiking activity, resembling previously reported oscillations in the rat subcortical visual system (Albrecht & Gabriel, 1994; Lewandowski *et al*., 2000; Szkudlarek *et al*., 2008; Chrobok *et al*., 2018). In Fig. 2C-E examples of rhythmically firing units in the dLGN, vLGN and OPN are presented. The mean frequency of observed oscillations ranged between 0.0022 and 0.018 Hz and did not differ between structures (Kruskal-Wallis test, K-W statistics = 0.8394, p = 0.6572; Fig. 2G). The proportion of infraslow oscillatory neurons was also statistically indistinguishable among these nuclei (28%, 22% and 30% in the dLGN, vLGN and OPN, respectively; Chi-square test, p = 0.3190, Chi-square = 2.285, df = 2, Fig. 3E). In the same dataset described in Fig. 2C-G we found that a fraction of neurons expressed fast narrowband oscillations in the beta/gamma range (20 – 100 Hz; Fig. 2H-J). A power spectra density (PSD) analysis and multimodal distribution of peaks at the level of ISI histograms of spontaneous spike activity revealed that 11% (dLGN), 15% (vLGN) and 33% (OPN) of all recorded units expressed such fast oscillations under dark-adapted condition. The frequency of these oscillations averaged at 33.43 ± 10.39 Hz (dLGN), 30.68 ± 6.16 Hz (vLGN) and 34.91 ± 5.09 Hz (OPN). In contrast to infra-slow oscillatory activity, the frequency and proportion of fast oscillating units differed between structures (Kruskal-Wallis test, p = 0.0271, K-W statistic = 7.213, Dunn’s multiple comparison test; Chi-square test, p < 0.0001, Chi-square = 30.29, df = 2; Fig. 2J, Fig. 3E).

**Fig. 2.**
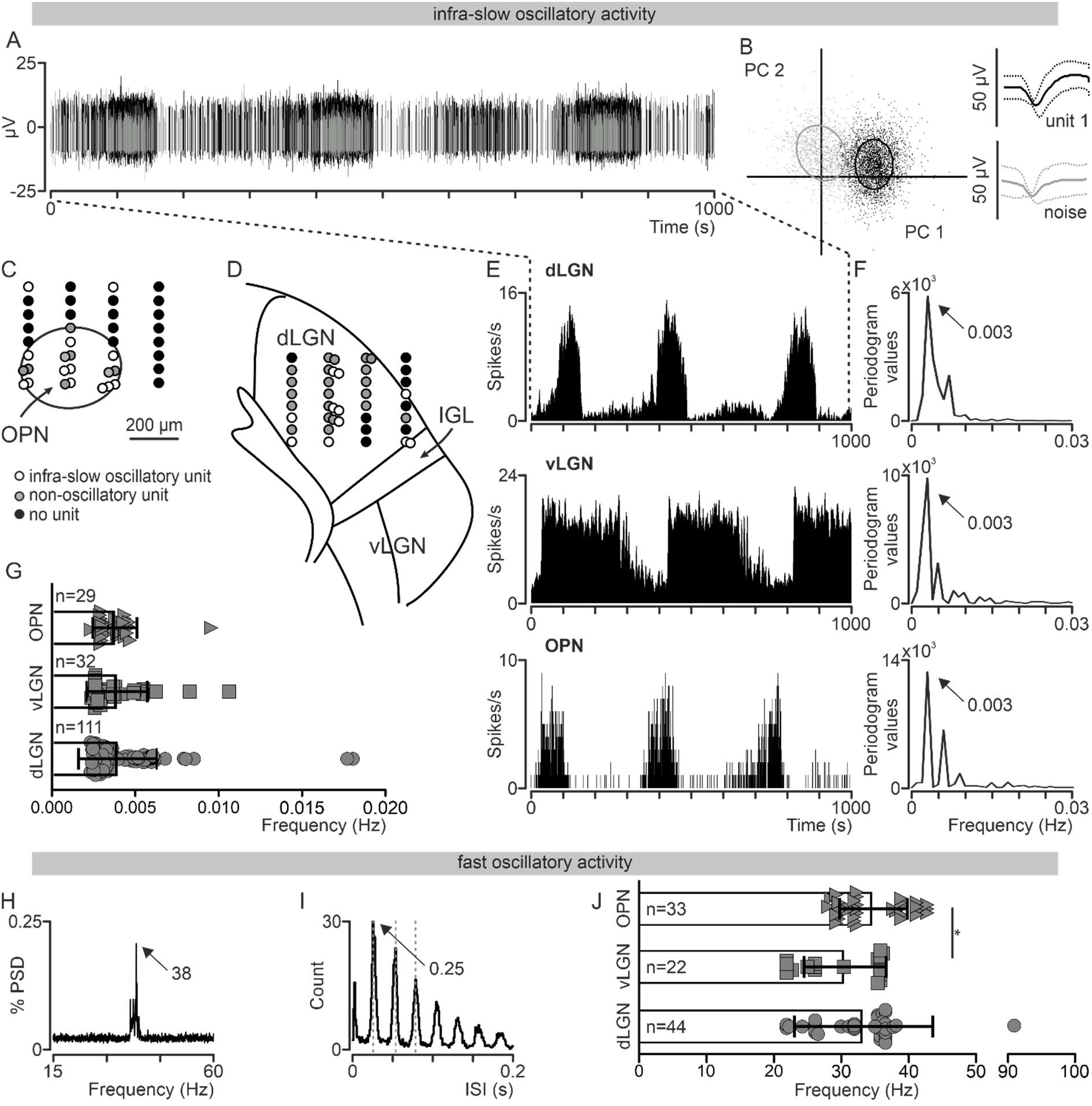
The presence of infra-slow and fast beta/gamma oscillatory activity in the mouse subcortical visual system. A) Representative trace recorded in the dLGN revealing infra-slow oscillatory activity in the dark-adapted spontaneous activity. B) Waveforms were clustered and sorted into units based on principal component analysis (PCA). A schematic representation of the placement of one 4 × 8 electrode (= 32 recording channels) showing the distribution of recorded units in the C) OPN and D) LGN and whether they were oscillatory (color coded). E) Representative firing rate histograms (bin size = 1 s) of spontaneous, dark-adapted activity of infra-slow oscillatory cells in the dLGN, vLGN and OPN (labeled above the histograms, not simultaneously recorded). F) FFT analysis performed for the recordings showed in E) indicating the infra-slow frequencies. G) The mean frequency of infra-slow oscillatory activity in the dLGN, vLGN and OPN does not differ between structures (Kruskal-Wallis test, K-W statistics = 0.8394, p = 0.6572). H) Fast beta/gamma oscillatory activity confirmed by PSD analysis. I) ISI histogram (bin size = 0.001 s) computed to confirm existence of harmonic modes in beta/gamma range. J) The mean frequency of fast (beta/gamma) oscillatory activity in the dLGN, vLGN and OPN differs between structures (Kruskal-Wallis test, K-W statistics = 7.213, p = 0.0271, Dunn’s multiple comparison test).

Based on that data (in total 640 cells from 30 recording sites from 22 wild type animals) we were able to describe four types of spontaneous activity (in darkness) in the sub-cortical visual system of visually intact mice: infra-slow or fast oscillatory activity alone; co-expression of both rhythms; and no oscillatory activity in the above ranges (Fig. 3A-E). Co-expression of infra-slow and fast oscillations were found in 5%, 6% and 15% of all dLGN, vLGN and OPN neurons, respectively (Chi-square test, p = 0.0022, Chi-square value = 12.24, df = 2, Fig. 3E). Importantly, all four types of neurons (infra-slow, fast, co-expressing both rhythms and none of them) were found simultaneously in 53% of the recording sessions indicating that their emergence cannot be trivially explained by global fluctuations in neuronal activity (Fig. 3F). In all the structures considered the expression of fast beta/gamma was limited to a minority of units. Simultaneous LFP recordings, reflecting population activity from all units, did not exhibit such narrowband oscillations (Fig. 3G-H).

**Fig. 3.**
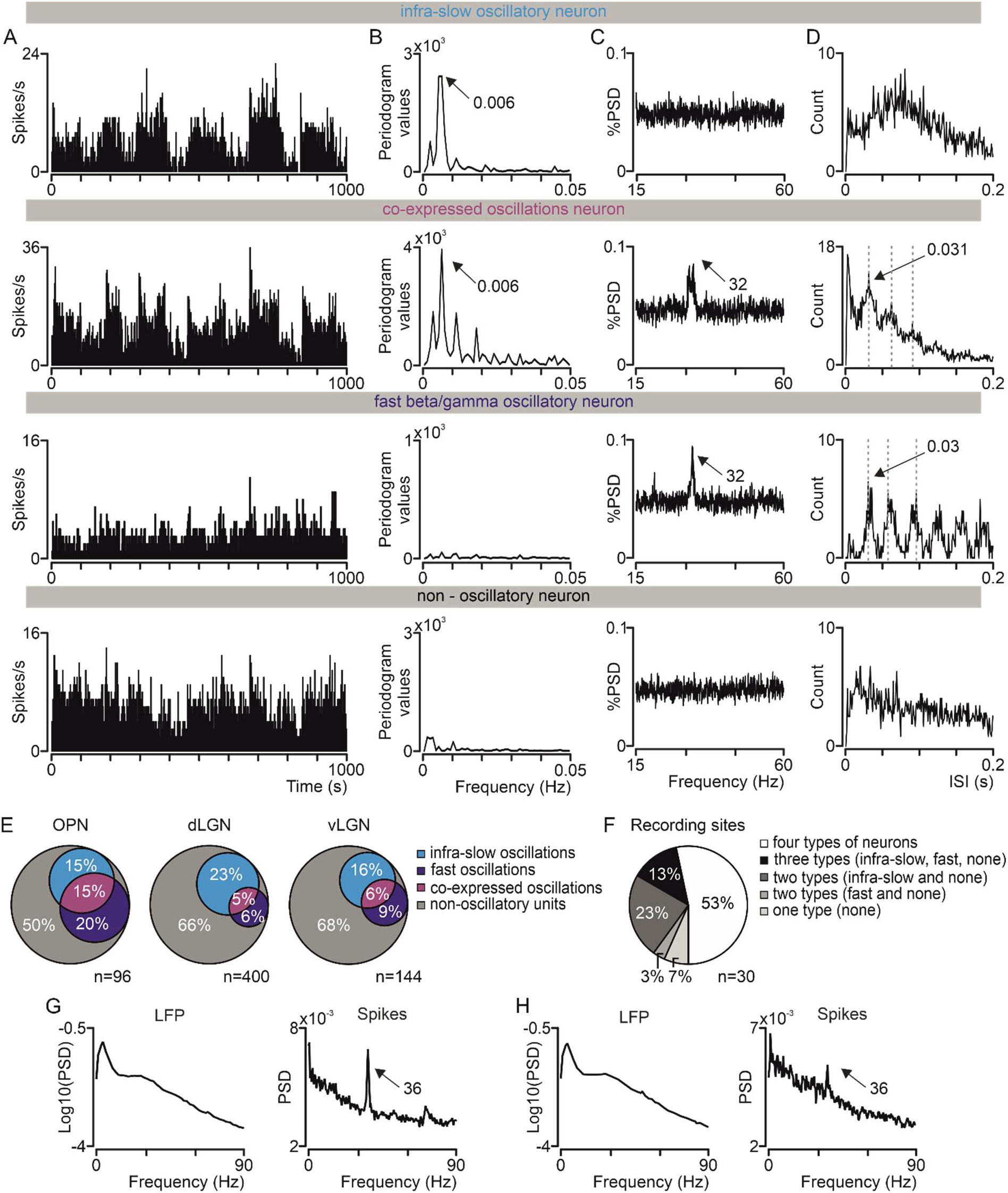
Types of spontaneous activity in the subcortical visual system. A) Representative firing rate histograms (bin size = 1 s) of spontaneous, dark-adapted activity of simultaneously recorded different types of oscillatory cells in the dLGN. B) FFT and C) PSD analysis were performed to classify cells as infra-slow, fast beta/gamma oscillatory, co-expressing both or none of the above frequencies. D) ISI histograms (bin size = 0.001 s) were computed to confirm existence of harmonic modes in beta/gamma range. E) Venn diagrams showing proportion of four different types of cells recorded in each structure. F) Pie chart summarizing proportion of recording sites (n = 30) in which infra-slow, fast, co-expressed oscillations or none of them were observed. G) PSD for LFPs (left) and for an individual unit (right) recorded from the same recording channel. Note that while a narrowband oscillation is clearly detectable from the spiking activity of the unit no clear narrowband peak can be observed in LFPs. H) Same as panel (G) but for a different recording channel during the same experimental session.

Firing rate of neurons expressing only infra-slow were significantly correlated between each other in all the structures investigated (except the OPN, Table 2 for *Opn1mwR* as revealed by the cross-correlation analysis (see Material and Methods section). Significant correlations were also observed in neurons co-expressing both infra-slow and fast oscillations (Table 2 for *Opn1mw^R^*). Moreover, these two populations were significantly correlated with each other (Table 2 for *Opn1mw^R^*) suggesting that a single infra-slow rhythm coordinates firing rate in both populations.

**Table 2:** Cross-correlation between infra-slow oscillatory neurons, neurons co-expressing infra-slow and fast oscillations and between these two populations. Data was analyzed by sign-test; all p-values < 0.0001 were set at 0; n – number of correlated pairs of neurons; bin size = 1 s. OPN – olivary pretectal nucleus, dLGN – dorsal lateral geniculate nucleus, vLGN – ventral lateral geniculate nucleus, *Opn1mw^R^* – visually intact mice, *Opn4^-/-^* – animals lacking melanopsin

| Structure | Infra-slow oscillations |  |  | Co-expressed infra-slow and fast oscillations |  |  | Between |  |  |
| --- | --- | --- | --- | --- | --- | --- | --- | --- | --- |
| *Opn1mw<sup>R</sup>* mice |  |  |  |  |  |  |  |  |  |
|  | n | p-value | r | n | p-value | r | n | p-value | r |
| dLGN | 420 | 0 | 0.238 | 27 | 0 | 0.370 | 82 | 0 | 0.187 |
| vLGN | 40 | 0 | 0.159 | 11 | 0.065 | 0.106 | 7 | 0.5 | -0.162 |
| OPN | 27 | 0.442 | 0.098 | 32 | 0 | 0.283 | 67 | 0.0002 | 0.118 |
| Pretectum | 45 | 0.036 | 0.086 | 4 | 0.125 | 0.127 | 7 | 1 | -0.0005 |
| *Opn4<sup>-/-</sup>* mice |  |  |  |  |  |  |  |  |  |
| dLGN | 89 | 0.0003 | 0.107 | 6 | 0.0312 | 0.310 | --- | --- | --- |
| vLGN | 37 | 0.099 | 0.096 | 5 | 0.375 | 0.213 | 21 | 1 | 0.079 |
| OPN | 15 | 0.001 | 0.090 | 6 | 0.031 | 0.306 | --- | --- | --- |
| Pretectum | 33 | 1 | 0.042 | --- | ---- |  | --- | --- | --- |

### Fast beta/gamma oscillations are modulated in the infra-slow range by diffuse co-expression of both rhythms at single unit level

We next aimed to investigate the possibility that infra-slow oscillations modulate fast beta/gamma oscillations. Specifically, we asked how and to what extent the phase of infra-slow oscillations could modulate the amplitude of fast beta/gamma oscillations.

We first investigated the possibility that this phenomenon could occur between two distinct neuronal populations, one expressing only infra-slow the other only fast beta/gamma. For each recording and brain structure, we calculated the phase of infra-slow and the amplitude of fast beta/gamma by separately pooling infra-slow and fast beta/gamma units and then by applying a frequency filter and a Hilbert transform to each population (see Methods for details). This analysis revealed that neurons expressing only fast oscillations were largely unaffected by the population-level infra-slow oscillations (Fig. 4A,B). We then asked whether this modulation could instead occur within the same population of neurons. Therefore we repeated the same analysis on populations co-expressing both infra-slow and fast beta gamma oscillations. We found that within these populations fast beta/gamma rhythms were strongly modulated according to the phase of infra-slow oscillations (Fig. 4D,E). These results were consistent across the whole dataset indicating that co-expression of both rhythms at the level of individual unit is required for infra-slow modulation of fast narrowband oscillations (Fig. 4C,F). Finally, we measured the extent to which the expressions of infra-slow and fast oscillations were coordinated at population level.

**Fig. 4.**
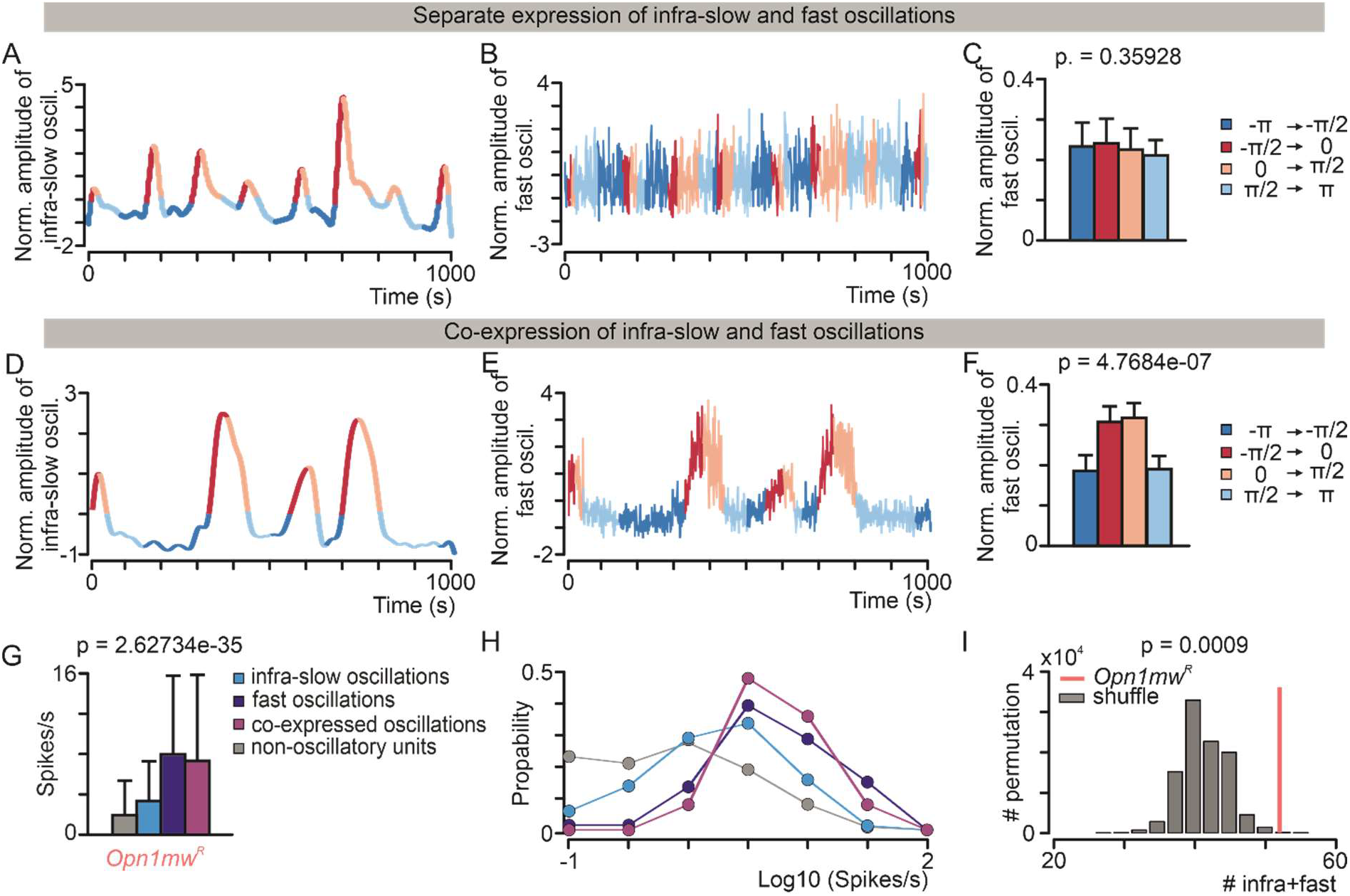
Infra-slow modulation of fast beta/gamma oscillations in the subcortical visual system of *Opn1mw^R^* mice. Representative example of simultaneously recorded activity from two separate pools of neurons. Firing rate activity has been filtered in the A) infra-slow and B) fast beta/gamma range and normalized as z-score. The color of the lines denote four different phases of the infra-slow rhythm according to the phase quadrants (dark blue: −π to −π/2 radians; dark red: −π/2 to 0 radians; dim red 0 to π/2 radians; dim blue: π/2 to π radians). C) A nonparametric sign-test comparison between fast beta/gamma amplitude recorded during the “red” vs the “blue” phases shows no significant phase modulation. Here the data are grouped across all experiments in which distinct neuronal pools expressing either infra-slow or fast beta/gamma were simultaneously identified. D) and E) are the same as A) and B), respectively, but for a representative example in which a pool of simultaneously recorded neurons co-expressed both infra-slow and fast oscillations. F) Same as panel C but for all experiments in which a population of neurons co-expressing both rhythms was identified. Within these populations the amplitude of fast beta/gamma is significantly modulated by the phase of infra-slow oscillations. G) Firing rates (mean ± SD) for all the four unit types described in **Figure 2A-E** (Kruskal-Wallis test, p=0, n = 822). H) Distribution of firing rates across the four unit types. I) Statistical comparison between the number of neurons co-expressing infra-slow and fast beta/gamma oscillations that we experimentally observed and the theoretical number predicted by the chance levels and based on the overall number of neurons expressing those two rhythms (either separately or jointly). The number of neurons in which infra-slow and fast beta/gamma are co-expressed (the red line) is larger than chance across our dataset. The null distribution (grey histogram) is obtained by using the permutation test as described in Material and Methods section. * < 0.05; ** < 0.01; *** < 0.001; **** < 0.0001

To address this question, we tested the possibility that co-expression of fast and infra-slow oscillations could simply occur by chance. The average firing rate of the four types of neurons were significantly different being highest for units not expressing fast beta/gamma (Fig. 4G; Kruskal-Wallis test, p = 0, n = 822). However the distributions of the four types as function of firing rate were largely overlapping (Fig. 4H) and this allowed us to perform a permutation test that preserved such distributions (see Permutation Test in Methods and Table 1 for details). Therefore we compared the observed number of units where the rhythms were co-expressed with the number we would expect under the null hypothesis that these two oscillations were independent (see Permutation Test in Methods and Table 1 for details). We asked whether a statistical dependence between infra-slow and fast oscillations expressing neurons would emerge from the whole. To do so we pooled together all neuronal groups recorded across structures. We found that the number of neurons co-expressing infra-slow and fast oscillations was larger than expected by chance, indicating that these rhythms tended to co-occur at the level of individual units (Fig. 4I).

### Outer and inner retina photoreception play different roles in the control of infra-slow modulation of fast beta/gamma oscillations

The data from visually intact mice thus suggest that infra-slow and fast oscillations are separable phenomena, which nonetheless co-occur more than expected by chance. Previous results indicate that expression of both infra-slow and fast beta/gamma oscillations depends both on outer and inner retina photoreception (respectively rods/cones and melanopsin) (Orlowska-Feuer *et al*., 2016a; Storchi *et al*., 2017; Chrobok *et al*., 2018). We now employed disruption of these two types of photoreceptors to further explore the inter-dependence of oscillations at high and infra-slow frequency. We repeated our initial experiments in a mouse model of aggressive retinal degeneration, lacking rod and cone photoreceptors at the time of recording *(rd/rd cl;* (Lucas & Foster, 1999; Lucas *et al*., 1999)), and in mice with a more subtle deficit in retinal function caused by loss of the inner retinal photopigment melanopsin *(Opn4^-/-^*). The detailed analysis of all recorded and analysed cells in all genotypes is presented in Table 3. Importantly, there were no differences in the mean firing rates between genotypes across the structures investigated (dLGN: Kruskal-Wallis test, Kruskal-Wallis statistics = 5.901, p = 0.0523; vLGN: Kruskal-Wallis test, Kruskal-Wallis statistics = 2.011, p = 0.3659; OPN: Kruskal-Wallis test, Kruskal-Wallis statistics = 5.740, p = 0.0567).

**Table 3.** Number and proportion of all recorded neurons. Infra-slow and fast beta/gamma oscillatory activities presented in different structures of the mice subcortical visual system and in different genotypes. Mean number of cells ± SD per recording site are presented in brackets. OPN – olivary pretectal nucleus, LGN – lateral geniculate nucleus, dLGN – dorsal lateral geniculate nucleus, vLGN – ventral lateral geniculate nucleus, *Opn1mw^R^* – visually intact mice, *rd/rd cl* – animals lacking rods and cones, *Opn4^-/-^* – animals lacking melanopsin

|  | No. of cells |  |  | No. of infra-slow oscillatory cells |  |  | No. of fast oscillatory cells |  |  |
| --- | --- | --- | --- | --- | --- | --- | --- | --- | --- |
| Genotype | *Opn1mw<sup>R</sup>* | *rd/rd cl* | *Opn4<sup>-/-</sup>* | *Opn1mw<sup>R</sup>* | *rd/rd cl* | *Opn4<sup>-/-</sup>* | *Opn1mw<sup>R</sup>* | *rd/rd cl* | *Opn4<sup>-/-</sup>* |
| dLGN | 400<br>(17 ± 8) | 258<br>(23 ± 10) | 340<br>(16 ± 8) | 111<br>(5 ± 6) | 107<br>(10 ± 12) | 53<br>(3 ± 2) | 44<br>(2 ± 3) | 5<br>(0.5 ± 0.8) | 23<br>(1 ± 2) |
| vLGN | 144<br>(8 ± 7) | 100<br>(11 ± 7) | 208<br>(10 ± 8) | 32<br>(2 ± 2) | 31<br>(3 ± 5) | 34<br>(2 ± 2) | 22<br>(2 ± 3) | 0 | 36<br>(2 ± 3) |
| OPN | 96<br>(14 ± 5) | 97<br>(10 ± 5) | 66<br>(9 ± 6) | 29<br>(4 ± 4) | 53<br>(5 ± 5) | 12<br>(2 ± 3) | 33<br>(5 ± 2) | 4<br>(0.4 ± 0.7) | 15<br>(2 ± 3) |
| Pretectum<br>(beyond<br>OPN) | 182<br>(23 ± 17) | 265<br>(22 ± 12) | 216<br>(27 ± 8) | 29<br>(4 ± 2) | 101<br>(8 ± 7) | 21<br>(3 ± 2) | 17<br>(2 ± 2) | 3<br>(0.25 ±<br>0.43) | 14<br>(2 ± 1) |
| Summary information |  |  |  |  |  |  |  |  |  |
| Genotype |  | *Opn1mw<sup>R</sup>* | *rd/rd cl* | *Opn4<sup>-/-</sup>* | Genotype |  | *Opn1mw<sup>R</sup>* | *rd/rd cl* | *Opn4<sup>-/-</sup>* |
| Number<br>of animals | Total | 22 | 17 | 22 | Number of<br>recorded/<br>analysed<br>cells | Total | 936/822 | 768/720 | 902/830 |
|  | OPN | 5 | 8 | 5 |  | OPN +<br>beyond | 300/278 | 364/362 | 312/282 |
|  | LGN | 17 | 9 | 17 |  | LGN | 636/544 | 404/358 | 590/548 |

In both groups of mice we found significant differences in the prevalence and the features of both infra-slow and fast beta/gamma oscillations compared with visually intact animals. The highest percentage of units expressing infra-slow oscillations was observed in *rd/rd cl* (41%, 31% and 56% for the dLGN, vLGN and OPN, respectively), followed by *Opn1mw^R^* (28%, 22% and 30%), and *Opn4^-/-^* mice (16%,16% and 18%,). Also the frequency of infra-slow oscillations was significantly different among genotypes. Specifically both *rd/rd cl* and *Opn4^-/-^* mice exhibited faster infra-slow oscillations compared with visually intact animals, however in the OPN that was only seen as a trend (dLGN: Kruskal-Wallis test, Kruskal-Wallis statistics = 37.53, p < 0.0001; vLGN: Kruskal-Wallis test, Kruskal-Wallis statistics = 11.05, p = 0.0040; OPN: Kruskal-Wallis test, Kruskal-Wallis statistics = 3.137, p = 0.2084; Fig. 5).

**Fig. 5.**
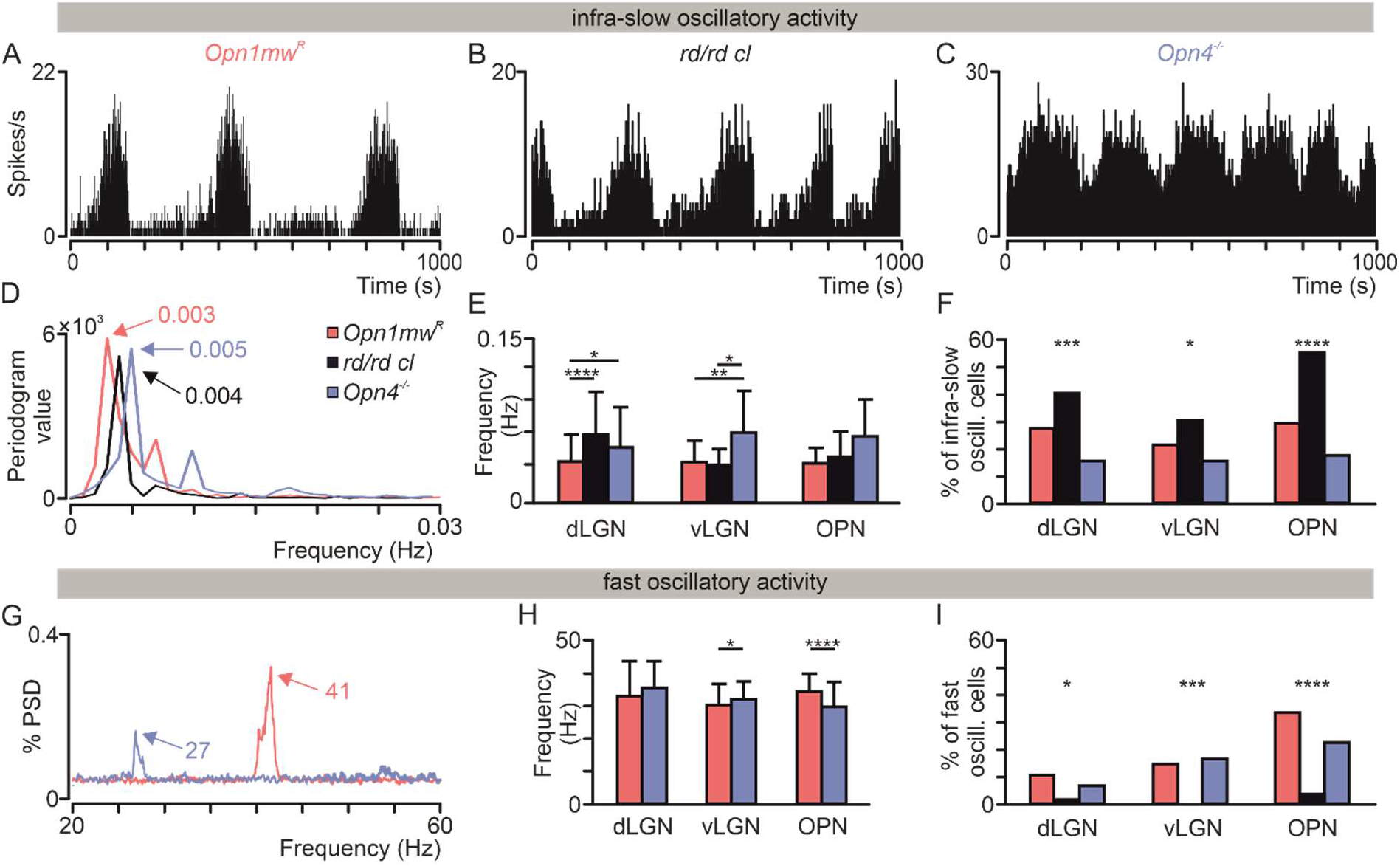
Inner and outer retina contribution to infra-slow and fast beta/gamma oscillatory rhythms emergence. Representative firing rate histograms (bin size = 1 s) of infra-slow oscillatory activity recorded within the dLGN of A) *Opn1mw^R^*, B) *rd/rd cl* and C) *Opn4^-/-^* mice. D) Peaks (color coded) correspond to the infra-slow frequency of oscillations presented on A – C, respectively. E) Mean frequency of recorded infra-slow oscillations differs between genotypes in the dLGN and vLGN, but not in the OPN (however the trend could be seen). F) The proportion of infra-slow oscillatory cells differs between the genotypes (Chi-square Test; dLGN: p = 0.0005, Chi-square statistic = 15.40, df = 2; vLGN: p = 0.0400, Chi-square statistic = 6.437, df = 2; OPN: p < 0.0001, Chi-square statistic = 33.32, df = 2). G) Normalized PSDs for two different OPN neurons recorded in *Opn1mw^R^* and *Opn4^-/-^* mice (colour coded). H) The mean frequency of fast beta/gamma oscillating neurons differs between *Opn1mw^R^* and *Opn4^-/-^* in the OPN and vLGN, but not dLGN. I) The proportion of fast beta/gamma oscillatory cells differs between the genotypes (Chi-square Test: dLGN: p = 0.0381, Chi-square statistic = 6.536, df = 2; vLGN: p = 0.0001, Chi-square statistic = 18.12, df = 2; OPN: p < 0.0001, Chi-square statistic = 28.44, df = 2;=). Please note, that fast oscillatory cells were not recorded at all in vLGN in *rd/rd cl* animals. Data in E analysed by Kruskal-Wallis test followed by Dunn’s multiple comparison test; data in H analysed by Mann Whitney. * < 0.05; ** < 0.01; *** < 0.001; **** < 0.0001

In *rd/rd cl* mice, in which almost half of the recorded units exhibited infra-slow oscillatory activity, fast oscillations were largely abolished (dLGN: 5 out of 258; vLGN: 0 out of 100, OPN: 4 out of 89). On the other hand, in *Opn4^-/-^* mice the percentage of units expressing fast oscillations was comparable with visually intact animals (7%, 17% and 23% in the dLGN, vLGN and OPN, Fig. 5I). However, the frequency of fast oscillations was altered in *Opn4^-/-^* mice in the OPN (lower in the *Opn4^-/-^* mice) and vLGN (higher in the *Opn4^-/-^* mice), but not in dLGN (Fig. 5H).

Since both rhythms were retained following melanopsin loss, we next asked whether this manipulation had altered the interaction between infra-slow and fast beta/gamma oscillations. Similarly to what we found in visually intact animals, neurons recorded from *Opn4^-/-^* mice could be classified into four distinct types on the base of their expression of one or either oscillatory activity. All four types of neurons were found in 28% of recording sites while in 13% of recordings we did not find any oscillatory activity (in total 614 units from 32 recording sites and 22 animals were analysed). Consistently with the *Opn1mw^R^* genotype, cross-correlation analysis revealed that in the dLGN firing rates were correlated within populations of neurons expressing only infra-slow or both rhythms (Table 2 for *Opn4^-/-^*). However, differently from visually intact animals, the firing rates were not correlated between these two populations (Table 2 for *Opn4^-/-^).* Infra-slow modulation of fast oscillations was observed, but as in visually intact mice, this only occurred in neurons expressing both rhythms (Fig. 6).

**Fig. 6.**
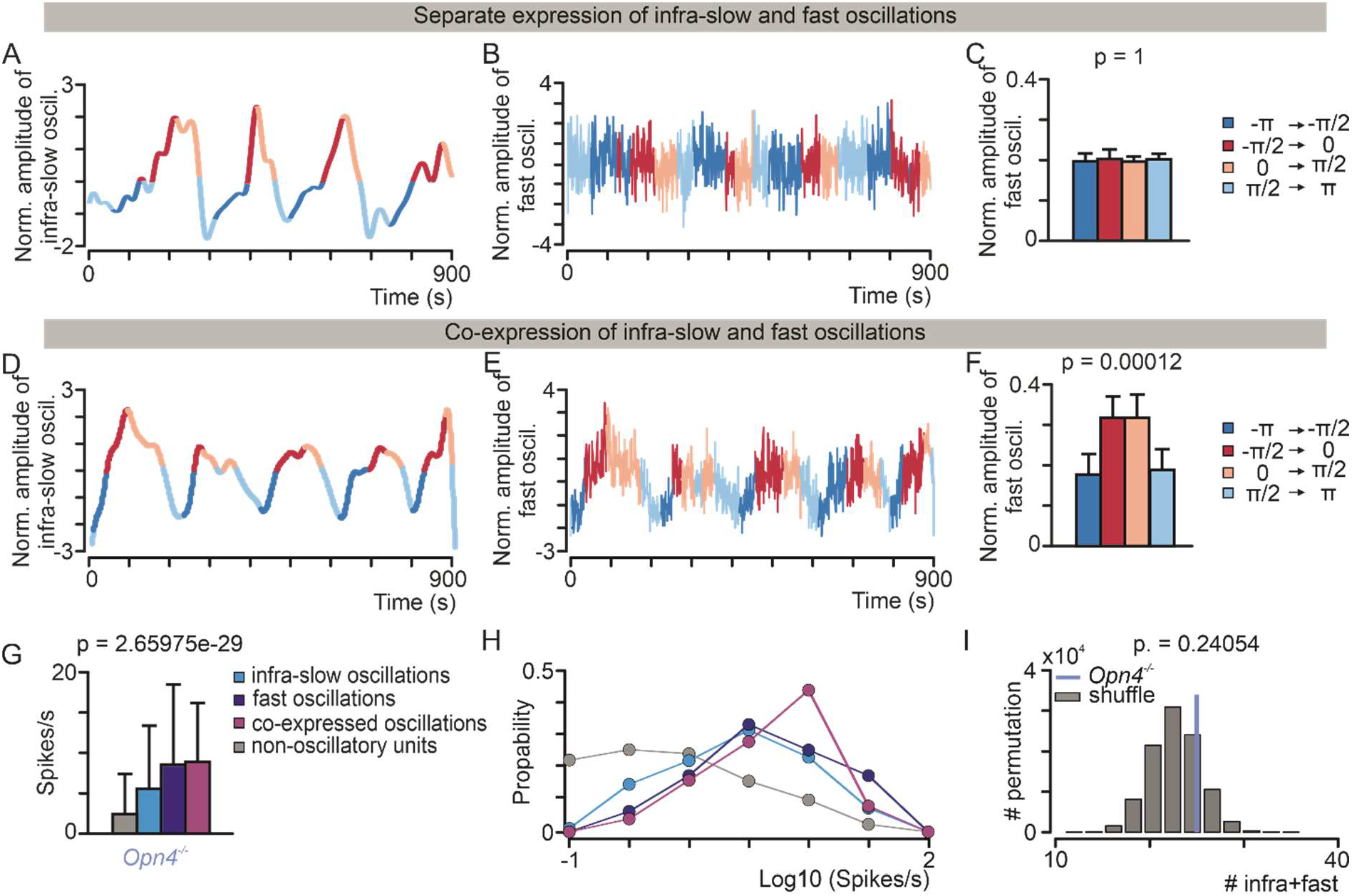
Infra-slow modulation of fast beta/gamma oscillations in the subcortical visual system of *Opn4^-/-^* mice. Same as Figure 4 but for the *Opn4^-/-^* dataset. * < 0.05; ** < 0.01; *** < 0.001; **** < 0.0001

Finally we asked whether the rate of co-expression of both oscillations was also preserved. We found that this was not the case since co-expression of infra-slow and fast beta/gamma oscillation was not significantly different from chance level in *Opn4^-/-^* mice (Fig. 6G-I). This result indicates that inner retinal photoreception, while not required for co-existence of both rhythms, is necessary for tying these rhythms together and enabling a diffuse modulation of fast beta/gamma oscillation in the infra-slow range.

## DISCUSSION

Infra-slow oscillations have been ubiquitously observed in the subcortical visual system, however their function remains poorly understood. Instead fast beta/gamma oscillations are involved in several types of visual processing that include inter-area communication along the visual pathway and feature binding (Gray *et al*., 1989; Fries, 2005). It was previously shown that gamma rhythm couples with slower rhythms expressed in different brain areas (Vanhatalo *et al*., 2004; Monto *et al*., 2008), thus we asked whether that is also the case with infra-slow oscillatory activity in the subcortical visual system.

We find that these two types of oscillation are distinct phenomena, but that they do co-occur more than expected by chance. Importantly, this co-occurrence allows the phase of infra-slow oscillations to modulate the amplitude of the fast. These findings reveal a potential mechanism by which infra-slow oscillations could regulate the visual flow – by modulating the amplitude of fast beta/gamma oscillations in the mouse visual system.

### Infra-slow oscillations are a common feature of subcortical visual systems

Oscillations with the period of minutes were described before in the visual system of adult rats (SCN: Miller & Fuller, 1992; dLGN: Albrecht & Gabriel, 1994; IGL: Lewandowski *et al*., 2000; OPN: Szkudlarek *et al*., 2008), cats (optic chiasm: Cavaggioni, 1968; retina: Rodieck and Smith, 1966; dLGN: Lorincz *et al*., 2009, Hughes *et al*., 2004), rabbits (visual cortex: Aladjalova, 1957), monkeys (visual cortex: Leopold et al., 2003; LGN: Cheong *et al*., 2011) and mice (prefrontal cortex, EEG recordings: Lecci *et al*., 2017; prefrontal cortex: Okun *et al*., 2019). Our study is the first to show the existence of infra-slow oscillatory rhythm in the mouse subcortical visual system, where we recorded them in the OPN and in both divisions of LGN. According to the present data, infra-slow oscillatory neurons constitute ~30% of all recorded units in dark adapted mice. The frequency of observed oscillations is slower (~0.005 Hz) than those of rats (~0.01 Hz), however, there is no doubt that the pattern of spike generation is the same (Fig. 2). Differences between these species in their frequency could have different potential origins. On one hand, it could be a genuine species difference in visual physiology. On the other, it may be connected to the fact that albino Wistar rats carry dysfunctions in the visual system (Bolles & Woods, 1964; Lund, 1964; Heiduschka & Schraermeyer, 2008; Nadal-Nicolás *et al*., 2012) while this study was performed in fully pigmented mice. Finally, it is possible that part of the variability could be simply explained by the small sample sizes used in the previous studies, where single channel recordings were employed as opposed to the higher-yield multichannel electrodes used in this study.

### Infra-slow and fast beta-gamma oscillations interact through co-expression of both rhythms at the level of single units

While co-existence of infra-slow and fast beta/gamma has been reported in the subcortical visual system (Tsuji *et al*., 2016; Chrobok *et al*., 2018), until now the extent to which they are independent events and whether they interact was unknown. Our data, obtained by unbiased recordings of all active neurons in the dark, reveal that while a subset of cells express only one type of oscillation, others express either both or none (Fig. 3).

In principle, in such population in which not all neurons expressing one type of oscillation also have the other, slow oscillations could still modulate fast beta/gamma oscillations by two types of circuitries. On the one hand, the population of infra-slow neurons could provide a modulation of beta/gamma oscillations, including neurons that did not themselves have infra-slow oscillations (Fig. 7A-F). Alternatively, such modulation could be restricted to neurons in which the two types of oscillations could interact within single neurons (Fig. 7H-J).

**Fig. 7.** Potential circuits mediating infra-slow modulation of fast beta/gamma oscillations. A-F) The amplitude of fast/beta gamma oscillations is modulated according to a distinct but functionally connected population expressing infra-slow oscillations. In this way coherence of oscillations in the beta/gamma range is modulated by the phase of infra-slow oscillations. A-C) Simulated firing pattern of a neuronal population expressing infra-slow oscillations (A) and magnified single units activity at different phase of the infra-slow oscillations (B). The population expresses changes in phase in the infra-slow range (C, top panel) but no fast beta/gamma oscillations (C, bottom panel). (D-F) A second neuronal population lacks infraslow modulation of firing rate (D) but exhibits epochs of regular fast beta/gamma oscillations (E, bottom) interleaved with epochs lacking such fast rhythms (E, top). The amplitude of fastbeta gamma oscillations is coupled to the phase of infra-slow activity (F). H) Simulated firing of a neuronal population expressing both infra-slow and fast beta/gamma oscillations. I) Magnified single units activity at different phase of the infra-slow oscillations. Clear beta/gamma oscillations can be observed during epochs of high firing rates. J) Within the same population the amplitude of fast-beta gamma oscillations is coupled to the phase of infra-slow activity.

Our analyses revealed that only neurons co-expressing both oscillations are responsible for mediating infra-slow modulation of fast beta/gamma oscillations (Fig. 4D-F). That is, when they co-occur the amplitude of fast modulations is modified according to the infra-slow rhythm. Importantly, the level of co-expression of these rhythms was higher than expected by two independent events suggesting that the interaction between infra-slow and fast beta/gamma occurs “by design” rather than simply by chance (Fig. 4G).

It is currently not possible to map the circuits displayed in Fig. 7 onto specific anatomical connections since the infra-slow modulation could be inherited by the retina or generated at the level of the subcortical visual system. Further *in vivo* studies performing simultaneous recordings from optic nerve and from neurons in subcortical visual system would be helpful to clarify the specific circuits involved.

### Changes to the retinal network alter appearance and interaction of infra-slow and fast oscillations

The retina has been previously implicated in the emergence of both infra-slow and fast oscillations in the subcortical brain, with intraocular injection of tetradotoxin abolishing both (Lewandowski & Błasiak, 2004; Szkudlarek *et al*., 2012; Chrobok *et al*., 2018). Moreover, selective blockade of phototransduction pathways (classic photoreceptors: rods and cones and melanopsin), as well as, retinal gap junction blockade, were shown to disrupt infra-slow oscillatory rhythm (Orlowska-Feuer *et al*., 2016a, 2016b). In all of the cited above data, intraocular injections were performed to assess retina contribution to the generation of infraslow oscillatory activity in the subcortical visual system. However such approach cannot distinguish between a driving effect, in which infra-slow oscillations in the retina and directly inherited by sub-cortical nuclei, and a gating effect, in which excitation by retina enables subcortical oscillations to emerge. Here, we studied rhythmicity in two mice strains in which the optic nerve was intact and active but in which retinal function was genetically disrupted. These were a model of aggressive photoreceptor degeneration, lacking rod+cone cells *(rd/rd cl* mice), and animals in which inner retina photoreception is ablated *(Opn4^-/-^* mice). As we were recording activity in the dark, the loss of photoreception in these manipulations is not itself expected to impact our findings, but secondary effects on the retinal circuitry could. Photoreceptors degeneration has knock-on effects on inner-retinal neurons and induces aberrant spontaneous ganglion cell activity in the form of ~10 Hz oscillations (Jones *et al*., 2003; Ryu *et al*., 2010; Menzler & Zeck, 2011; Trenholm & Awatramani, 2015), while melanopsin knockout has been reported to impact aspects of retinal development (Renna *et al*., 2011; Rao *et al*., 2013; Kirkby & Feller, 2013).

Infra-slow oscillations were retained in all genotypes. In fact, they were most prevalent in the *rd/rd cl* (see Table 3), consistent with the simulated effect of reduced retinal input (Zierenberg *et al*., 2018). Both genetic modifications altered their frequency, and this effect was replicated across all visual centers, consistent with the view that these infra-slow oscillations share a common mechanistic origin.

The situation with respect to fast beta/gamma oscillations was quite different, with this behavior essentially absent in *rd/rd cl* mice. This dramatic effect highlights the importance of the retinal activity for this rhythm. Since fast beta/gamma oscillations are typically network phenomena (Buzsáki & Wang, 2012), that could be because the retinal circuits generate these oscillations (Kenyon *et al*., 2003); that interplay between central and retinal circuits is required to produce this rhythmicity; or that the change in retinal input to the brain alters the activity of central circuits generating this rhythmicity.

Perhaps the most interesting alteration in oscillatory activity in transgenic mice was the effective decoupling of infra-slow and fast oscillations in melanopsin knockout animals. While in intact animals these two rhythms co-occurred in individual neurons more often than chance, this property was lost in the *Opn4^-/-^* genotype. Although we can only speculate as to the change in retinal circuitry responsible for this disruption, it does reinforce the conclusion that the cooccurrence of these oscillations in visually intact animals is an active process. It has been shown that vascular patterning of the retina is dependent upon intact melanopsin photoreception, since this process is disrupted in both dark reared visually intact animals and in animals lacking melanopsin photoreception (Rao *et al*., 2013). Since vascular patterning affect development of neuronal networks in the retina, this could also affect the coupling between infra-slow and fast narrowband rhythms. This result also provides a new potential mechanism via which this inner retinal photoreceptor typically associated with sub-conscious reflex responses to light could contribute to conventional visual function. While it has been previously shown that the psychophysical threshold of visual sensitivity oscillates in the infra-slow range (Thoss *et al*., 1998) the neuronal mechanism underlying this phenomenon is currently unknown. The reported coupling between infra-slow fluctuation and fast beta/gamma oscillations, the latter being involved in visual processing, points towards such a potential mechanism.

### Limitations of the study

Since the presented results were obtained from urethane-anesthetized mice an open question is to what extent those results generalize to awake behaving animals. While a number of studies have reported infra-slow and betta/gamma oscillations in various species under different experimental conditions e.g. anesthetics (such as barbiturates, isoflurane) and recording procedures (LFPs, EEG, *in vitro* and *in vivo* neuronal recordings) (Albrecht *et al*., 1998; Filippov & Frolov, 2005; Hughes *et al*., 2011; Storchi *et al*., 2017) some studies have also reported such rhythms in the dLGN and primary visual cortex in behaving animals (Albrecht *et al*., 1998; Filippov & Frolov, 2005; Saleem *et al*., 2017; Storchi *et al*., 2017). The frequency of both types of oscillations was higher in freely moving compared to anaesthetized animals (Albrecht *et al*., 1998; Storchi *et al*., 2017) and such difference could be due to the specific action of the anesthetics on neurotransmission systems (Hara & Harris, 2002; Wu *et al*., 2004; Hemmings *et al*., 2005). Coupling between infra-slow and fast beta/gamma oscillations has also been recorded in awake cortical recordings in human (Monto *et al*., 2008) but, to the best of our knowledge, never in the subcortical visual structures where our recordings were performed.

All recordings in the present study were performed under dark-adapted condition. It has been previously shown that both, infra-slow and beta/gamma oscillatory neurons are modulated by light and visual stimuli. Photopic backgrounds increase mean firing rate but not oscillatory frequency in neurons expressing infra-slow oscillations (Szkudlarek *et al*., 2012; Chrobok *et al*., 2018, however see also (Albrecht *et al*., 1998)). In neurons expressing narrowband gamma oscillations both amplitude and frequency of such oscillations are enhanced by steady light (Saleem *et al*., 2017; Storchi *et al*., 2017). Those previous results indicate that bright lights strongly increase the expression of fast narrowband oscillations and, consistently with this notion, expression of fast beta/gamma oscillations under dark-adapted conditions is limited to a minority of neurons. While individual units can still express narrowband oscillations such oscillations were not detectable in LFP recordings. This result is in accordance with previous studies indicating that the coupling between LFPs and spikes only becomes apparent with high synchrony at population level (Denker *et al*., 2011; Storchi *et al*., 2012). During spontaneous activity, synchrony is strongly reduced in comparison with evoked sensory responses and, as a result, coupling between and between LFP and spikes and between LFPs from different subcortical areas is also reduced (Storchi *et al*., 2012; Zippo *et al*., 2013).

### A potential role for the interaction between infra-slow and fast beta/gamma oscillations

In considering the potential function of these two types of oscillation it is important to remember that they have very different spatiotemporal topology. Infra-slow fluctuations are sustained events that can last minutes/hours and are correlated between different brain areas (Szkudlarek *et al*., 2008; Chrobok *et al*., 2018). Infra-slow oscillations have also been closely associated with Resting State Networks (RSNs) observed in human fMRI recordings (Damoiseaux *et al*., 2006; De Luca *et al*., 2006; He *et al*., 2008), and most likely reflect the underlying structural connectivity (Fox & Raichle, 2007; Honey *et al*., 2007). Fast beta/gamma oscillations typically are short-lasting events that can arise locally and transiently synchronize distant brain areas (Buzsáki, 2006). Those oscillations can be evoked by sensory stimulation (Gray *et al*., 1989; Cardin *et al*., 2009; Storchi *et al*., 2017) or by cognitive tasks (Fries, 2005) but can also emerge spontaneously (Tsuji *et al*., 2016; Chrobok *et al*., 2018; Foik *et al*., 2018). Thus, their spatiotemporal topology is more flexible and need not necessarily reflect the underlying structural connectivity. Accordingly, the interaction between infra-slow and fast beta/gamma could represent a “balancing act” between the opposing needs for transient “on-demand” flexibility – served by fast beta/gamma oscillation – and long-term structural and dynamical stability – served by infra-slow oscillations.

Viewed from this perspective, the strong modulation of fast beta/gamma amplitude by infraslow oscillations observed here could represent a mechanism to oppose runaway plasticity (Abbott & Nelson, 2000) and promote the stability of visual signaling. This control would then be significantly reduced in *Opn4^-/-^* animals since these two rhythms are independently expressed within the subcortical visual networks. Consistent with this possibility a previous work showed that light modulation of fast beta/gamma oscillations is substantially reduced in *Opn4^-/-^* animals (Storchi *et al*., 2017). Thus, although fast beta/gamma could be observed in the dLGN of *Opn4^-/-^* animals, these oscillations were less controllable by the visual stimulus. This lack in controllability could be due to the lack of the stabilizing action of infra-slow oscillations.

## ADDITIONAL INFORMATION

### Data availability statement

The data that support the findings of this study are openly available in github at https://github.com/RStorchi/InfraSlow.

### Conflict of interest statement

The authors declare no competing interests.

### Funding

The work of POF was supported by the Bekker Programme implemented by the Polish National Agency for Academic Exchange (NAWA). RS was funded by National Centre for Replacement Refinement and Reduction of Animals in Research (NC3Rs) via a David Sainsbury Fellowship (NC/P001505/1). RJL was funded by The Welcome Trust (Investigator Award 210684/Z/18/Z) and The European Research Council (268970).

### Author contributions

POF, HJS, RJL, RS designed research; POF, AEA, TMB performed research; RS contributed analytic tools; POF, HJS, RS analyzed data; POF, RJL, RS interpreted the results; POF, RJL, RS wrote the paper. All authors have approved the final version of the manuscript and agreed to be accountable for all aspects of the work in ensuring that questions related to the accuracy or integrity of any part of the work are appropriately investigated and resolved. All persons designated as authors qualify for authorship and all those who qualify for authorship are listed.

## Notes

https://github.com/RStorchi/InfraSlow

### Competing Interest Statement

The authors have declared no competing interest.

